# Is Protein Quantification and Physical Normalization Always Necessary in Proteomics?

**DOI:** 10.64898/2026.02.13.705808

**Authors:** Alex Zelter, Michael Riffle, Gennifer E. Merrihew, Batool Mutawe, Aaron Maurais, Han-Yin Yang, Jamie L. Inman, Susan E. Celniker, Jian-Hua Mao, Kenneth H. Wan, Antoine M. Snijders, Christine C Wu, Michael J. MacCoss

## Abstract

Dogma suggests protein quantification is a pre-requisite to LC-MS/MS based proteomics studies. Such quantification allows a standardized ratio of sample to digestion enzyme and enables physical normalization of protein digest loaded onto the mass spectrometer for analysis. Most proteomics studies include these steps. However, there are significant costs in time, money and experimental complexity, associated with performing protein quantification and physical normalization for every sample, especially for larger studies. Proteomics data analysis pipelines typically include computational normalization strategies to compensate for unavoidable systematic biases. These strategies also have the potential to compensate for avoidable variation such as omitting sample amount normalization. Here we investigate the effects of either physically normalizing the amount of protein for each individual sample or leaving it unnormalized. Our results show the relationship between increased protein amount variation in sample input, and the variance of quantified relative abundances of peptides and proteins output after data analysis. The experiments presented here suggest that protein quantification and physical normalization steps can be omitted from some quantitative proteomic experiments without incurring an unacceptable increase in measurement variability after computational normalization has been applied. This work will enable important time and cost saving optimizations to be made to many proteomics workflows.

## Introduction

Protein quantification has historically been considered an essential pre-requisite for liquid chromatography tandem mass spectrometry (LC-MS/MS) based proteomics studies.^1,2^ Such quantification is performed on the proteins to be analyzed (e.g. tissue lysate) prior to digestion into peptides. Quantification of the amount of protein in the samples allows the correct ratio of sample to digestion enzyme to be calculated for each sample and additionally enables normalization of the quantity of protein digest loaded onto the mass spectrometer for analysis. The dogma is that this is necessary, especially for quantitative proteomic studies. Some reasons typically given as to why this physical normalization is necessary are improved digest reproducibility and improved quantification.^1,2^

The reasons outlined above are valid, and most proteomics studies currently perform protein quantification and physical normalization steps prior to sample digestion and analysis. In a world of unlimited resources, these steps would indeed be carried out for every sample. However, it is important to recognize that there are significant costs, in both time, money and experimental complexity, associated with performing such quantification and standardization for every sample. This is especially true for large scale studies involving hundreds or even thousands of individual samples.

For specific sample types processed using a standardized protocol, the range of protein concentrations typically liberated can often be well defined with a limited number of initial measurements. With this data in hand, two important questions can be raised: (1) What is the cost to my experimental results of accepting that range of protein concentrations versus the cost of quantifying and physically normalizing the amount of protein in each individual sample to be analyzed? (2) Which cost is most acceptable given the questions I am trying to answer and the resources I have available to answer them?

Regardless of whether the amount of protein digested and analyzed is physically normalized or not, proteomics data analysis pipelines include computational data normalization strategies designed to compensate for systematic biases that are unavoidably introduced during sample processing and data collection.^3–5^ These strategies also have the potential to compensate for avoidable variation introduced during sample processing, such as omitting physical normalization.

In the current work we investigate the effects of either physically normalizing the amount of protein in each individual sample or leaving it unnormalized and accepting the resulting range of protein sample amounts in the hope that it can be sufficiently compensated for using computational normalization strategies applied after the mass spectrometry data is collected.

Our results show the relationship between increased protein amount variation in sample input and the variance of the quantified relative abundances of peptides and proteins output after data analysis. The data presented suggest that, for many experiments, protein quantification and physical normalization steps can be omitted from quantitative proteomic workflows with only a modest reduction in sensitivity and/or increase in measurement variability after computational normalization is applied, while still yielding data that are sufficient to address the intended experimental questions. This work will enable important time and cost saving optimizations to be made to many proteomics workflows.

## Methods

### MatTek Tissue Culture and Sample Irradiation

MatTek EpidermFT full-thickness skin tissues (MatTek Corporation, EFT-400) were maintained according to the manufacturer’s recommendations. Tissues were cultured at 37°C with 5% CO₂ in serum-free, phenol red-free maintenance medium (MatTek EFT-400-MM-PRF), with 2.5 mL of pre-warmed medium replaced daily. On day 0, immediately prior to irradiation, 300 µL of conditioned medium was collected from each well for baseline Lactate Dehydrogenase (LDH) assessment and replaced with fresh medium. LDH release was quantified using the Cytotoxicity Detection Kit (Roche, 11644793001) following the manufacturer’s instructions. A standard curve was generated using a dilution series of the provided LDH standard solution. Conditioned media were collected from individual MatTek EpidermFT transwells, and 100 µL per sample was transferred to a 96-well plate. Absorbance was measured at 490 nm using a SpectraMax iD3 plate reader. Per manufacturer guidance, wells with LDH readings ≥ 2.0 were excluded from analysis. LDH assessment was subsequently performed every three days to monitor tissue health.

Irradiations were performed using an XRAD-320 cabinet irradiator (Precision X-Ray) equipped with a 0.5-mm copper (Cu) filter plate, a rotating platform stage, an adjustable table, and a fully open collimator aperture. Samples were positioned at a source-to-sample distance of 50 cm. The irradiator was operated at 300 kV and 10 mA, delivering dose at a rate of 1.3 Gy/min. Sample doses can be found in the MatTek metadata is available here: https://panoramaweb.org/sample-normalization.url.

### MatTek Sample Production

For the MatTek physically normalized (PN) versus not physically normalized (NPN) experiment, two batches of MatTek EpiDermFT skin tissue were cultured and irradiated as described above. Samples were harvested at 1-, 6-, and 14-days post-irradiation. Individual tissues were removed from transwells under sterile conditions and placed on a cutting board within a Class II biosafety cabinet. An 8-mm biopsy punch (Integra Miltex, 33-37) was used to obtain full-thickness biopsies, which were then quartered using a No. 10 scalpel (Integra Miltex, 4-410). Each quarter was placed in a pre-labeled Protein LoBind tube (Eppendorf, 022431081), flash-frozen in liquid nitrogen, and stored at –80°C until further processing.

MatTek lysates were produced by thawing each sample on ice, immersing in 150 µL of lysis buffer (defined above) and sonicating for 10 seconds at level 3 followed by 30 seconds at level 4. Once homogenized, the samples were immediately stored over dry ice.

Sonicated MatTek samples were centrifuged at 16,873 x g for 10 minutes and the supernatant was transferred to a new Eppendorf tube. A BCA assay was performed on all samples using Invitrogen’s Pierce BCA Protein Assay Kit according to the manufacturer’s instructions (cat no. 23227).

For physically normalized (PN) samples a variable volume of lysate plus a variable volume of diluent (lysis buffer) was combined to result in each sample having a final volume of 100 µL containing 50 µg total protein. For not physically normalized samples (NPN) a fixed volume of lysate (30 µL) was combined with a fixed volume of diluent (70 µL) resulting in a final volume of 100 µL per sample. This fixed volume was determined based on average protein concentrations from previous experiments on different MatTek samples prepared in a similar way. BCA results from the current sample set were not used in determining the fixed lysate volume used here.

### Mouse Pelt Sample Collection Procedure

C57BL/6 and BALB/c mice were anesthetized with isoflurane, and body weight was recorded. Mice were euthanized by cervical dislocation under isoflurane anesthesia. Following euthanasia, mice were shaved and pelts were dissected using a ventral midline incision. The fourth mammary gland was removed from the inner surface, and the exposed area was wiped with sterile gauze. To maintain orientation, pelts were pinned and flipped to expose the dorsal side. The central dorsal region was sampled by collecting eight 5-mm skin punches using disposable biopsy punches (Integra Miltex, catalog #33-35). Biopsies were removed with sterile forceps, transferred into sterile 1.5 mL microcentrifuge tubes, and flash-frozen in liquid nitrogen.

### Mouse Pelt Lysate Pool Production

Pelt punches were placed in LoBind Eppendorf tubes and immersed in 250 µL lysis buffer consisting of 100 mM Tris, pH 8.5/2% SDS + HALT Protease and Phosphatase Inhibitor Single-Use Cocktail (Thermo Scientific, Catalog number PI78442). Samples were sonicated using a Misonix S-400 / Microtip sonicator set at 34% amplitude (6.75 kHz) for 10 seconds followed by 4 cycles of 40% amplitude (8 kHz) for 30 seconds then rest for 30 seconds with total sonication time between 2 and 4 minutes.

Pelt lysates were individually centrifuged in a microfuge for 10 min at 1,000 x g and the supernatant was transferred to a new tube. A BCA assay was performed according to the manufacturer’s instructions (Invitrogen’s Pierce BCA Protein Assay Kit, part number 23227) to determine the protein concentration of each individual lysate. Equal volumes of all lysates were combined to make a pelt lysate pool and a separate BCA was done to determine the concentration of the final pool.

A variable volume of pelt lysate pool plus a variable volume of diluent (100 mM tris pH 8.5, 2% SDS) was added to a KingFisher 96 well plate (Thermo Fisher Scientific catalogue no. 95040450) such that the target total protein amount was achieved for each digest amount tested. Five replicates of each digest amount depicted in Figure 1b were prepared. The amounts were: 8.33, 16.7, 33.3, 50, 66.7, 83.3 and 100 µg of protein.

**Figure 1:**
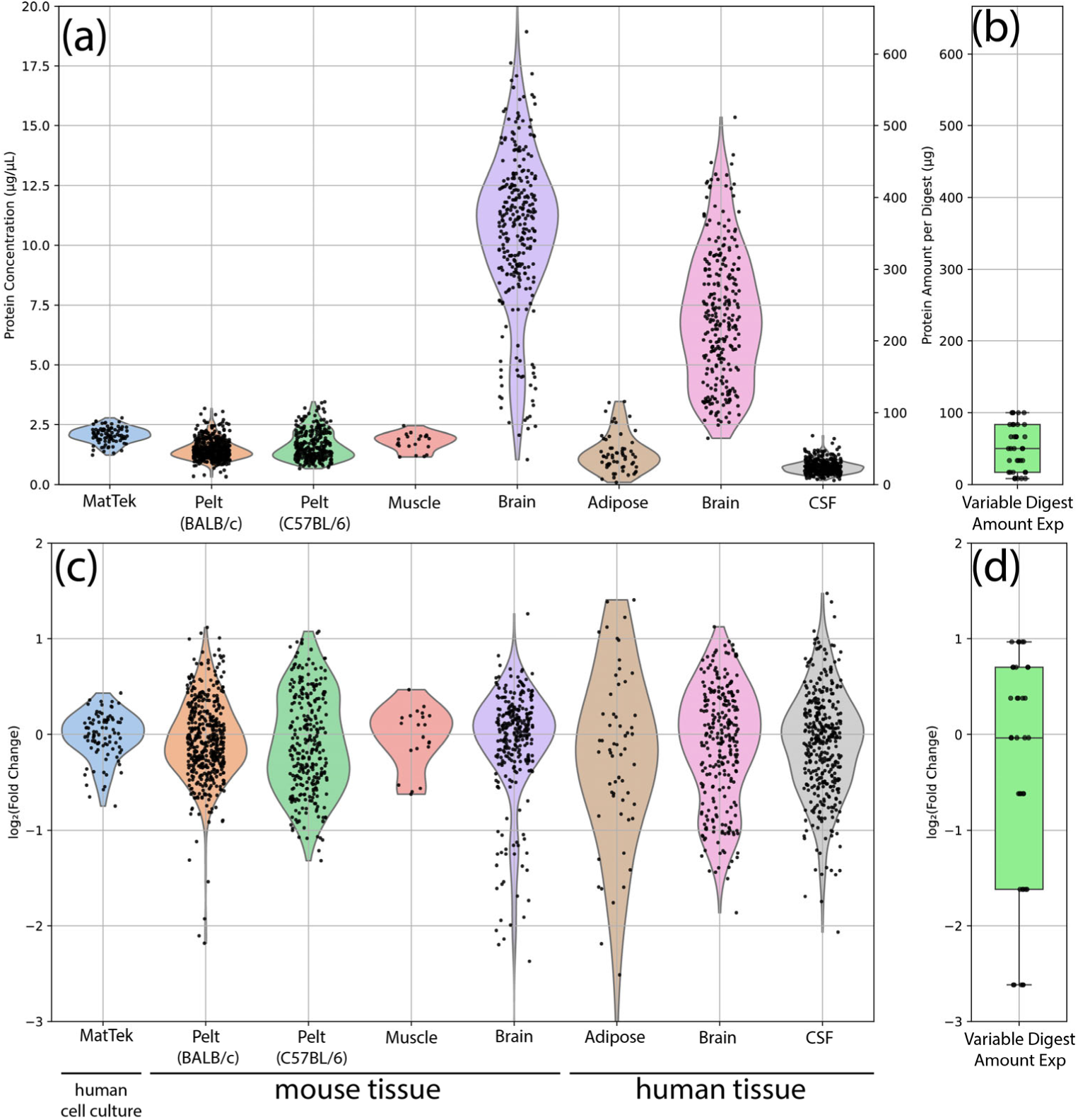
Different lysis protocols and sample types yield a range of protein lysate concentrations. To model this variability, we designed a variable digest amount experiment. (a) Protein concentrations (µg/µL, left axis) were measured by BCA. These values were also converted to “protein amount per digest” (right axis) assuming a fixed digest volume of 33.3 µL and a nominal lysate concentration of 1.5 µg/µL to show how deviations from the assumed concentration would alter the actual protein input for an experiment targeting 50 µg protein per digest. (b) A variable digest amount experiment was performed using defined amounts of a mouse pelt lysate mixture. The amounts were chosen to span a range around the target of 50 µg per digest, reflecting the variability in lysate concentrations observed in (a). (c) and (d) The data from (a) and (b) are shown as log_2_(fold change). For each sample, fold change was calculated relative to the mean concentration of its sample type.

### Protein Concentration Measurements for Figure 1

The protein concentrations of MatTek tissue culture samples along with C57BL/6 and BALB/c mouse pelt samples were measured by BCA as part of the current work as described above. Mouse muscle protein concentrations were from Campbell et al., 2019.^6^ Mouse brain protein concentrations were from Saul et al., 2024.^7^ Human adipose, brain and cerebrospinal fluid (CSF) protein concentrations were from Zelter et al., 2026,^8^ Merrihew et al., 2023^9^ and Tsantilas et al., 2024,^10^ respectively.

### Protein Digestion

Samples for mass spectrometry analysis were prepared by protein aggregation capture (PAC).^1,2^

**For the variable digest amount experiment** using mouse pelt pools each well in the 96 well plate described above contained 80 µL of lysate. Each well subsequently received 20 µL of an enolase (Sigma, cat no. E6126)/TCEP (Invitrogen, Bond-Breaker TCEP Solution, cat no. 77720) mixture dissolved in 50 mM Tris pH 8.5. Final TCEP concentration was 10 mM and each well contained 240 ng of enolase, which was added as a process control.^3^ A lid was placed on the plate (Corning, cat no. 3099) and samples were reduced for 1 hour at 37°C in an Eppendorf thermomixer shaking at 650 rpm. Alkylation was initiated by adding 3 µL 500 mM iodoacetamide followed by incubation in the dark for 30 minutes at room temperature. The reaction was quenched by addition of 2 µL 500 mM DTT. To each well, 12.5 µL of MagResyn Hydroxyl particles (ReSyn Biosciences, cat no. MR-HYX002) were added and protein was aggregated to the beads by addition of 470 µL of acetonitrile (80% final concentration).

The beads were cycled through 5 KingFisher deep well wash plates containing 1 mL of wash solution per well (3 plates containing 95% acetonitrile followed by 2 plates containing 70% ethanol) with each wash step taking 2.5 minutes in a KingFisher Apex (Thermofisher Scientific). Protein-containing beads were released into 150 µL 50 mM tris, pH 8.5 containing 2.5 µg trypsin (Pierce, cat no. PI90058) per well. This amount of trypsin results in a 1:20 trypsin to substrate ratio assuming 50 µg sample protein per well, which is our typical target protein amount for this type of digest. In the current study protein amount was not always 50 µg thus the trypsin to sample ratio will also vary in such samples. Digestion was performed at 47°C for 1 hour in the KingFisher.

After digestion, 120 µL per sample was added to a 1.5 mL LoBind Eppendorf tube containing 6 µL 10% trifluoroacetic acid (TFA). Acidified peptides were spun in a bench top centrifuge at maximum speed for 10 minutes and 50 µL of sample was added to a LoBind Eppendorf tube containing 5 µL of 500 fmol/µL Pierce Peptide Retention Time Calibration Mixture (PRTC) (Thermo Scientific, cat no. PI-88321) dissolved in 0.1% TFA, added as a process control.^3^ From each digest, 5 µL of peptides + PRTCs were pooled into a single Eppendorf tube for generation of a chromatogram library^4^ as described below in the Data Processing section.

**For the MatTek physically normalized (PN) versus not physically normalized (NPN) experiment,** each well contained 100 µL of lysate. Each well was brought up to 200 µL by addition of 100 µL 100 mM tris pH 8.5 containing enolase and TCEP such that the final amount of enolase per well was 800 ng and TCEP was 10 mM final concentration. Samples were reduced for one hour as described above, followed by alkylation by addition of 10 µL 300 mM iodoacetamide and incubation in the dark for 30 minutes at room temperature. The reduction reaction was quenched by addition of 10 µL 300 mM DTT. To each well, 12.5 µL of MagResyn Hydroxyl particles were added and protein was aggregated to the beads by addition of 767 µL of acetonitrile (77% final concentration).

The beads were washed, and samples were digested as described above for the defined sample concentration testing experiment. After digestion, 100 µL of sample was moved to a 96 well autosampler plate (Thermo Fisher Scientific, catalogue number 60180-P207B) containing 7 µL of 10% TFA per well and mixed by pipetting. 50 µL of acidified sample was moved to a similar 96 well autosampler plate containing 5 µL of 500 fmol/µL PRTC per well and mixed by pipetting. From each digest, 5 µL of peptides + PRTCs of all PN samples were pooled into a single Eppendorf tube and 5 µL of peptides + PRTCs of all NPN samples were pooled into a second Eppendorf tube for generation of a chromatogram libraries^4^ as described below in the Data Processing section. Plates were then sealed using sealing tape (Thermo Fisher Scientific catalogue number 60180-M146) and stored at −80°C prior to analysis.

### Mass Spectrometry

Digested protein samples were analyzed by data independent acquisition-mass spectrometry (DIA-MS). Each sample contained enolase and PRTCs, which were added as process controls as described above.^10^ In all cases, a Thermo Vanquish Neo UHPLC System was used to load 6 µL of sample using a “trap and elute” workflow with a Thermo Scientific PepMap Neo Trap Cartridge (cat no. 174500) and a Bruker PepSep C18 15 cm x 150 µm, 1.9 µm column (cat no. 1893471). The HPLC column outlet was attached to a 5 cm x 20 µm ID Sharp Singularity (Fossil Ion Tech) tapered emitter mounted in a custom-built microspray source maintained at 45°C. The following acetonitrile gradient was used to elute peptides from the column: (1) 0-0.7 mins; 3-5% B; flow 1.3 µL/min; (2) 0.7-1 mins; 5-5.5% B; flow 1.3 µL/min; (3) 1-57 mins; 5.5-35% B; flow 0.8 µL/min; (4) 57-57.5 mins; 35-50% B; flow 1.3 µL/min; (5) 57.5-58 mins; 50-99% B; flow 1.3 µL/min; (6) 58-60 mins; 99% B; flow 1.3 µL/min. Buffer A was 0.1% formic acid in water. Buffer B was 0.1% formic acid, 80% acetonitrile and 20% water.

A Thermo Fisher Scientific Orbitrap Eclipse was used to collect mass spectrometry data in DIA mode using two methods. The first method was performed on each all individual sample and used 12 m/z overlapping precursor isolation windows^11,12^ with the following parameters. The Orbitrap resolving power for MS1 and MS/MS were 30,000 at *m/z* 200. Each cycle consisted of 1 MS scan followed by 75 MS/MS scans. MS scan range was *m/z* 395–1,005. Automatic gain control was set to standard for MS scans and to a normalized value of 800% for MS/MS scans. MS/MS spectra were acquired using overlapping isolation windows of 12 *m/z* with 75 MS/MS scans per cycle and a normalized collision energy of 27 assuming charge state 3. Cycle 1 spanned 406.4347 *m/z* through 994.7021 *m/z* in 12 *m/z* isolation windows. Cycle 2 spanned 400.4319 *m/z* through 1000.7048 *m/z* in 12 *m/z* windows. All spectra were collected in centroid mode.

The second method enabled creation of an on-column chromatogram library to improve analysis of the 12 m/z DIA data collected from individual samples. This library was generated by performing gas-phase fractionation on a pooled sample prepared by combining all individual samples from the experiment.^13,14^ The chromatogram library was generated from six LC-MS/MS runs, each spanning a different portion of the full *m/z* range used for the individual sample analyses. The six scan ranges were 395–505, 495–605, 595–705, 695–805, 795–905, and 895–1005 *m/z*. Data were acquired using 4 *m/z* staggered isolation windows, while all other settings, particularly the HPLC separation conditions, were kept identical to those used for the individual sample runs acquired with 12 *m/z* precursor isolation windows.

### Data Processing and Normalization

For the variable input experiment, data was processed using an automated Nextflow^15^ workflow, nf-skyline-dia-ms version a59bee31ae7997ce1e70f6fc10aa6c9d765c374b (available at: https://nf-teirex-dia.readthedocs.io/en/latest/), which performed the following general tasks: a) Demultiplexing^11,12^ of raw MS data and conversion to mzML files. This was done using ProteoWizard’s msConvert^16^ version 3.0.24172 using the following arguments: --mzML --zlib --ignoreUnknownInstrumentError --filter "peakPicking true 1-" --64 --filter "demultiplex optimization=overlap_only" –simAsSpectra; b) Generation of an on-column chromatogram library as previously described^14^ using narrow-window DIA data analyzed with EncyclopeDIA (version 2.12.30) plus a Prosit^17^ predicted spectra library based on the UniProt mouse FASTA; c) Quantification of wide-window data using EncyclopeDIA version 2.12.30 plus the on-column chromatogram library generated in the previous step; d) Generation of extracted ion chromatograms and export of peak areas using Skyline;^18^ e) Conversion of Skyline reports to unnormalized and normalized precursor and protein level reports was done using an inhouse produced Nextflow workflow available at: https://github.com/uw-maccosslab/nf-dia-batch-correction. This workflow applied Median Deviation (MD) normalization or total ion current (TIC) normalization at the precursor (peptide) or protein level.^9^ Normalization was performed across the entire sample set and both unnormalized and normalized precursor and protein values were output as tsv files, which were used in downstream analyses. Original Skyline documents and tsv files are provided on Panorama Public^19^ here: https://panoramaweb.org/sample-normalization.url, with values provided as log_2_(area+1), where area is the sum of the total area of all MS/MS transitions for a given peptide (peptide level) or all measured peptides in a given protein (protein level). Data analysis and figure plotting was performed on the resulting reports using the inhouse python code available here: https://github.com/uw-maccosslab/manuscript-sample-normalization.

Data for the MatTek experiments were processed similarly, except that an instrument-specific, optimized, spectral library in dlib format was generated using file the UniProt reviewed human proteome (UP000005640) plus DIA-NN^20^ version 1.8.1 and Carafe version 0.0.1 as previously described.^21^ This was done using a Nextflow workflow, nf-carafe-ai-ms version e6eab37fd6, available at: https://nf-carafe-ai-ms.readthedocs.io/en/latest/ and the resulting spectra library was used in place of the Prosit mouse library used in step (b) above.

### Logistic regression-based classification

For each of the four combinations of physical normalization (applied or not) and computational normalization (applied or not), we trained a logistic regression classifier to distinguish whether a sample was exposed to any radiation dose. Computational normalization consisted of median normalization, in which each sample was divided by its median value and then scaled by the mean of medians across all samples, followed by ComBat^22^ batch correction using the plate variable from the metadata (available here: https://panoramaweb.org/sample-normalization.url) as the batch covariate. Classifiers were implemented using scikit-learn’s LogisticRegression,^23^ with ElasticNet regularization^24^ specified by l1_ratio = 1 and C = 0.5. Model performance was evaluated as the area under the receiver operating characteristic (ROC) curve (AUC), estimated using 5-fold stratified cross-validation (RepeatedStratifiedKFold) repeated five times.

### Data Availability

Raw MS data, sample metadata and quantified peak areas associated with the current work are available on Panorama Public^19^ here: https://panoramaweb.org/sample-normalization.url and will be assigned a ProteomeXchange^25^ ID upon publication.

### Code availability

MSConvert is available from https://proteowizard.sourceforge.io/. Carafe, EncyclopeDIA, and Skyline are available from https://github.com/Noble-Lab/Carafe, https://bitbucket.org/searleb/encyclopedia/, and https://skyline.ms/skyline.url, respectively.

The Nextflow workflows used for spectral library generation, database searching, quantification, and normalization of the mass spectrometry data presented in this manuscript are publicly available. Spectral libraries generated using Carafe and DIA-NN were produced using nf-carafe-ai-ms (https://nf-carafe-ai-ms.readthedocs.io/en/latest/). On-column chromatogram libraries followed by quantification using EncyclopeDIA and Skyline were generated using nf-teirex-dia (https://nf-teirex-dia.readthedocs.io/en/latest/). Peptide- and protein-level normalization was performed using nf-dia-batch-correction (https://github.com/uw-maccosslab/nf-dia-batch-correction). All in-house code used for analysis of the data presented in this work is available at https://github.com/uw-maccosslab/manuscript-sample-normalization.

## Results and Discussion

### Protein Concentration Range Across Sample Types

The typical protein concentration from a given sample type depends on the protocol. For example, the same sample lysed in twice the volume of buffer will have a lower protein concentration. Likewise, twice the amount of sample lysed in the same volume will have a higher protein concentration.

The protein concentration for a given sample type produced using a fixed protocol depends on the sample. After running a limited number of BCA assays for a specific sample type–protocol combination, one can select a fixed lysate volume that results in the desired protein amount per digest. If individual samples are not physically normalized, variation in protein per digest will then reflect the natural variation among samples of that type.

We previously performed BCA-based protein quantification on various sample types as part of our laboratory’s historical standard operating procedure for proteomic analyses. Figure 1 shows the range of protein concentrations in lysates from a selection of sample types.

Large differences in protein concentrations from different sample types were observed (Figure 1a). Brain tissue samples gave the highest concentrations (median 6.9 µg/µL) while cerebrospinal fluid (CSF) produced the lowest (median 1.2 µg/µL).

While different protocol-tissue type combinations may yield different typical protein concentrations, it is the natural variation in concentrations among samples of a given type that is of most concern if samples are not to be physically normalized. To better visualize this variation, the data in Figure 1a was re-plotted to show the log_2_ fold change of each concentration measurement, calculated relative to the mean concentration of its sample type. We observed from about +1 to −1.5 log_2_ fold change across all the sample types (Figure 1c). Variation was typically greater in samples taken from human subjects versus tissue from experimental animals (Figure 1c, human versus mouse samples) and was lowest in tissue culture samples (Figure 1c, MatTek samples).

### Effect of Protein Input Variation on Measurement Precision

We tested how variation in protein amount per digest influences the precision of peptide and protein measurements reported after data analysis. To do this, specific amounts of a single mouse pelt pool (Figure 1b, Table 1) were digested and analyzed by data independent analysis (DIA).^26–28^ This method allows untargeted quantitation of extracted precursor and product ion chromatograms (XICs) for all detected peptides within the sampled m/z range (in this case 400-1000) without a list of pre-specified protein or peptide targets.

**Table 1:** Amount of mouse pelt lysate protein digested per replicate type.

| Sample ID | Amount of protein per digest ($\mu\text{g}$) | Amount of protein in 6 $\mu\text{L}$ injection ($\mu\text{g}$) |
| --- | --- | --- |
| 0.25 | 8.33 | 0.33 |
| 0.5 | 16.7 | 0.67 |
| 1.0 | 33.3 | 1.33 |
| **1.5** | **50.0** | **2** |
| 2.0 | 66.7 | 2.67 |
| 2.5 | 83.3 | 3.33 |
| 3.0 | 100 | 4 |

Five replicates were prepared for each sample digest amount. The series was designed such that the middle value, 50 µg, represents the “target” amount of protein typically digested per sample in our laboratory’s standard digestion protocol. The variation in protein amounts tested (Figure 1b and d) were based on the range measured in real samples (Figure 1a and c). For each replicate, 6 µL of digest was injected into the mass spectrometer. Additionally, similar replicates were prepared, and their protein concentration was re-measured by BCA without digestion.

The correlation between the calculated protein load - based on the initial BCA-measured pelt lysate pool concentration - and the protein amount measured by BCA after dilution was excellent (r² = 0.99) across the measured range (Figure 2a). However, the absolute protein amounts differed between calculated and measured values. This discrepancy is common and reflects inter-assay variability in BCA-based protein quantification when measurements are performed on different days or by different operators.

**Figure 2:**
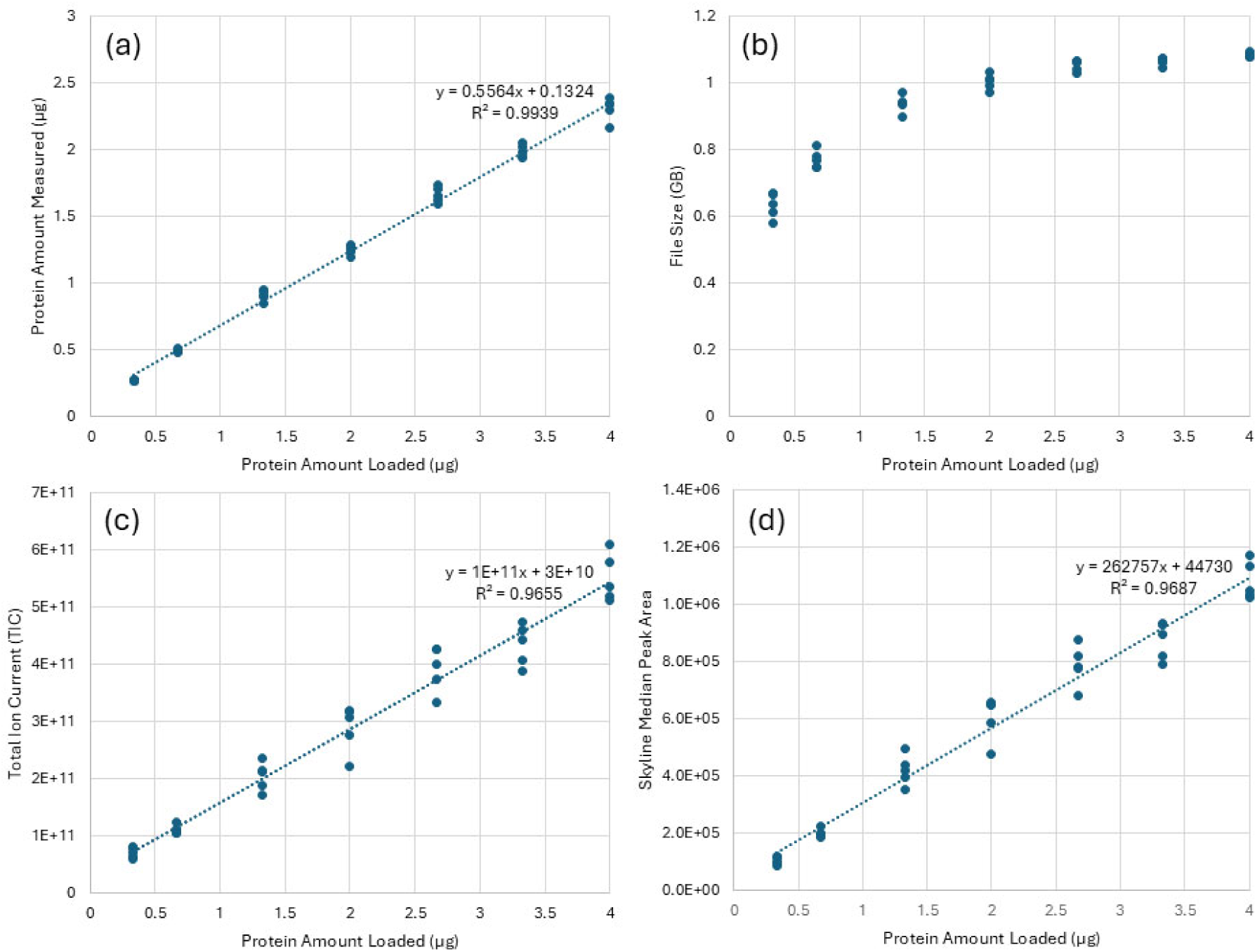
The relationship between the amount of protein loaded on to a mass spectrometer, or into a BCA assay, and signal. The four panels represent: (a) BCA measured protein amount, (b) mass spectrometer raw data file size, (c) mass spectrometer measured total ion current (TIC) and (d) mass spectrometer measured median peak area. In each case the “Protein Amount Loaded” was calculated based on the original BCA measured protein concentration of the mouse pelt pool and the known dilution and volume of each sample produced.

The raw data file size for each sample analyzed increased with increasing amounts of protein injected; however, the relationship was not linear (Figure 2b). The relationship between the amount of protein injected and the MS1 total ion current (TIC) was linear (r² = 0.97) across the range tested (Figure 2c). Similarly, the median peak area of all peptides quantified in each sample was linear (r² = 0.97) across the tested range (Figure 2d).

The median peak area increased with increasing amounts of protein injected. This is expected as the more analyte one injects the higher the signal should be. It also highlights a justification, in quantitative proteomics, to physically normalize the amount of protein per digest and subsequently the amount of protein injected into the mass spectrometer. This justification states that if the amount of protein injected into the mass spectrometer is not controlled the resulting quantitative values will primarily reflect differences in sample loading rather than true differences in protein abundance relative to the total proteome, thereby obscuring the biological changes that most proteomics experiments aim to measure. This justification is true but can be alleviated by computational normalization.

Most proteomics data analysis pipelines include a computational normalization step to compensate for unavoidable systematic biases. Common examples of computational normalization strategies include TIC and median normalization^9,29^. Given that both TIC and median peak area have an excellent linear relationship with the amount of protein injected into the mass spectrometer, it is unsurprising that these metrics have been found effective in normalizing differences between sample loading amounts.

In addition to compensating for unavoidable systematic biases, computational normalization strategies have the potential to correct for avoidable biases, allowing recovery of the relative abundance measurements of interest, even when combining data gathered on samples loaded in different amounts. To test the extent to which this is possible, we quantified all detectable peptides in the above mouse lysate digest dilutions. Precursor and protein values were calculated for all replicates individually (n=5 at each protein amount). Computationally unnormalized peak areas showed an increase with increasing protein amount loaded. Both TIC and median normalization removed this relationship (Figure 3).

**Figure 3:**
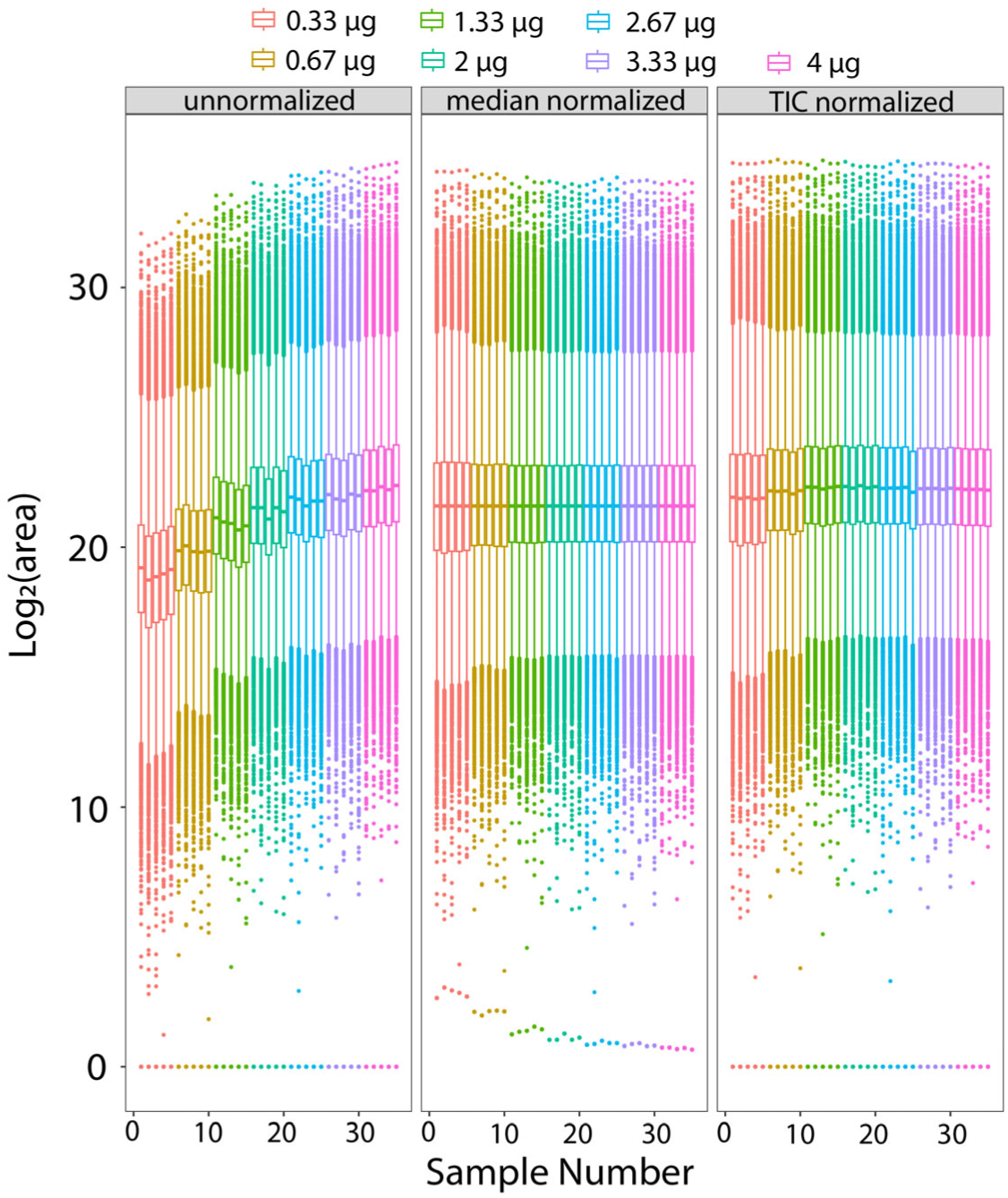
Relationship between protein amount loaded and Log_2_ peak area of quantified precursors with and without computational normalization. A pooled protein sample was diluted to five concentrations (n=5 for each concentration), digested, and analyzed by LC–MS/MS. The same injection volume was used for all samples, resulting in different total amounts of protein loaded onto the mass spectrometer. Precursor peak areas were quantified for each sample and box plots are shown of all samples in order of increasing protein concentration with no normalization, median normalization and TIC normalization.

Replicates for each sample digest amount were then placed into groups of different protein amount ranges (Figure 4, Protein Amount Groups, inset - bottom right) and the distribution of the coefficient of variation for all peptides and proteins in each group were plotted (Figure 4). This was done using no computational normalization, TIC normalization and median normalization.

**Figure 4:**
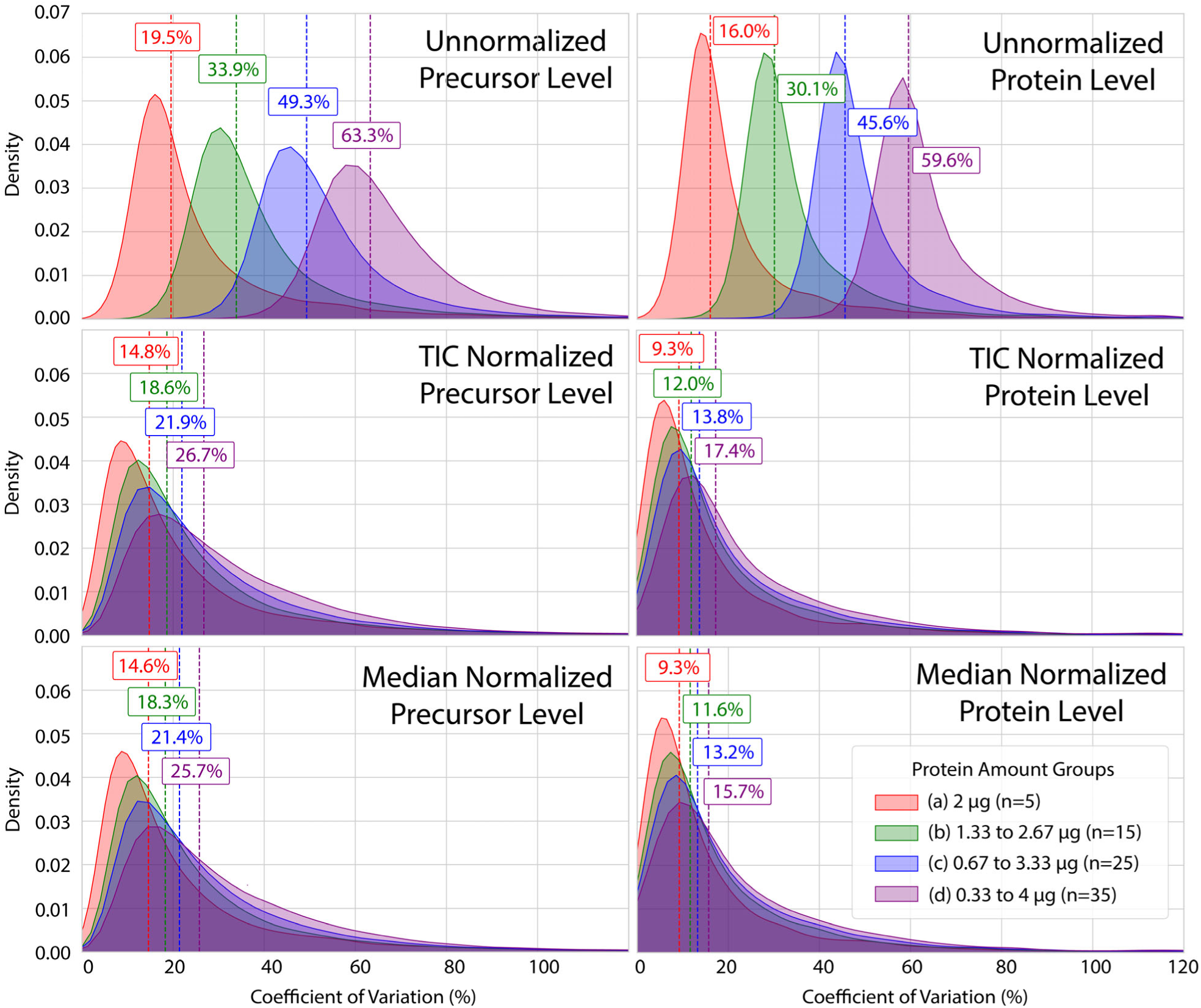
Relationship between protein amount variation and measurement precision when combining samples across different loading groups. Density plots show CV distributions computed after progressively combining data from an expanding range of protein-loading groups (bottom right inset key). CVs are lowest when only a single protein amount group is included (all replicates contain the same injected amount) and increase as additional higher and lower protein amounts are incorporated due to systematic changes in signal with analyte load. TIC and median normalization substantially reduce this variance increase, partially compensating for differences in protein amount across injections.

Combining computationally unnormalized data from replicates containing only a single protein amount per digest resulted in a median CV of 19.5%. This rose to 63.3% when combining data from replicates spanning the full range of protein digest amounts tested (Figure 4, top left panel). This increase highlights the legitimate concern that leads to laboratories performing physical normalization in their workflows. TIC normalization compressed this increase such that the median distribution of variance across the different groups increased from 14.8% to 26.7%. Median normalization performed similarly.

A similar analysis was performed after combining precursor peak areas into protein-level peak areas. At the protein level, the median distribution of variance across the different protein amount groups increased from 16% to 59.6% (Figure 4, top right). TIC and median normalization compressed this range such that replicates containing only a single protein amount per digest resulted in a median CV of 9.3% increasing to 17.4% or 15.7% across all groups for TIC and median normalized data, respectively.

Computational normalization is therefore capable of reducing a substantial amount of the variance in quantified precursor and protein measurements associated with increased variance in the amount of protein injected into the mass spectrometer. The key question is whether this compensation is sufficient to allow omitting physical protein amount normalization from quantitative proteomic experimental workflows while still enabling the aims of an experiment to be satisfied.

### Hypothesis Testing Without Protein Amount Normalization

We assessed whether quantitative proteomics experiments can yield meaningful results without physical protein amount normalization using a set of 96 MatTek EpiDermFT skin tissue samples. These samples were exposed to varying amounts of ionizing radiation prior to harvesting (MatTek metadata is available here: https://panoramaweb.org/sample-normalization.url). Protein lysates were produced using a set protocol and digests were set up in one of two ways: 1) A set volume of each lysate plus diluent was added to each digest with no prior knowledge of sample protein concentration. The volume was based on prior BCA-based protein concentration measurements made from historical samples prepared by our lab using the same protocol. The volume chosen targeted approximately 50 µg per digest. These samples are refered to as not physically normalized (NPN). 2) A BCA assay was performed on all sample lysates and a variable volume of lysate plus a variable volume of diluent was added to each digest. This resulted in a fixed amount of protein in every digest. These are referred to as physically normalized (PN) samples.

For all samples a fixed volume of 6 µL was injected into the mass spectrometer for analysis. This volume corresponded to 2 µg total protein for the physically normalized samples and an unknown amount of protein for the not physically normalized samples. Samples were analyzed using DIA, and all detectable peptides were quantified. The resulting peak areas were used to train a logistic regression classifier to distinguish whether each sample had been exposed to radiation or not.

Each of the two data sets (physically normalized or not physically normalized) was used either with or without applying computational normalization resulting in 4 different combinations. For each combination of physical normalization (applied or not) and computational normalization (applied or not), we trained a separate classifier. Model performance was evaluated as the area under the receiver operating characteristic (ROC) curve (AUC). An AUC of 1 represents a perfect classifier and an AUC of 0.5 indicates a predictive performance equivalent to random chance.

On computationally unnormalized data, the classifier reached an AUC of 0.83 without physical normalization. With physically normalized samples, performance improved to an AUC of 0.95 (Figure 5). On computationally normalized data, however, the classifier was able to reach a near-perfect score of 0.95 without physical normalization. Physical and computational normalization combined boosted performance by 4% to an AUC of 0.99. These data show that in this scenario an effective classifier, able to distinguish irradiated from non-irradiated samples, could be trained without measuring sample lysate protein concentrations and without physically normalizing the amount of protein digested for each sample. Omitting physical normalization did, however, increase dependence on effective computational normalization to achieve the experimental aims.

**Figure 5:**
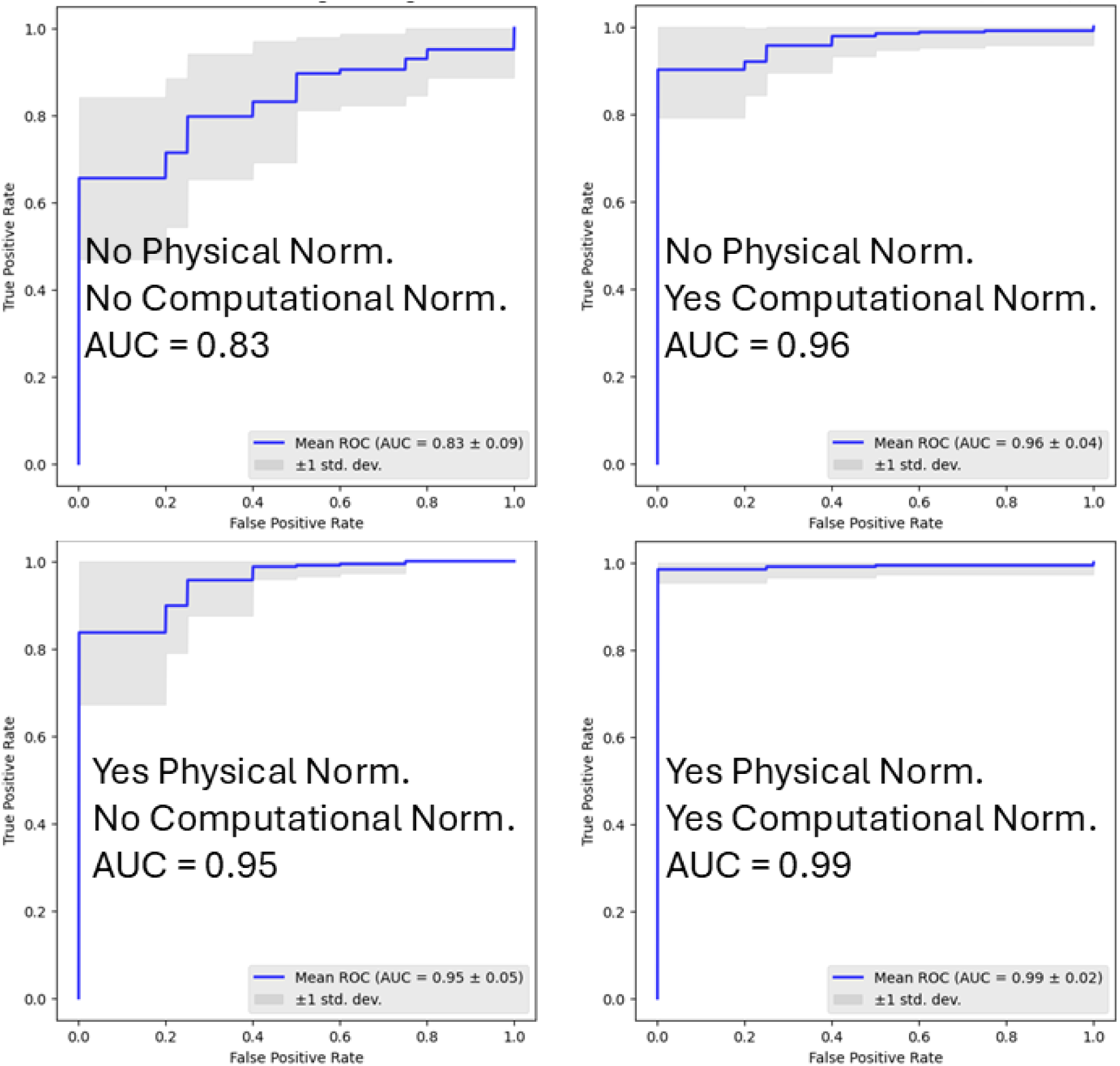
Quantitative proteomics experiments performed with and without physical and/or computational normalization. ROC curves resulting from logistic regression classification of radiation exposure from physically or computationally normalized or unnormalized data.

## Conclusions

Current thinking suggests that proteomics studies require physical sample normalization. This involves measuring the protein content of each sample and adjusting it to a consistent amount before digestion and analysis. Such physical normalization comes at a cost in time, money and experimental complexity. This cost increases with the size of the experiment.

In the current work we show that although physical normalization of samples improves the precision of quantified peptide and protein values, computational normalization can largely compensate for omitting this step. We perform side-by-side experiments with or without physical normalization, which show that real-world experimental objectives can be met without physical normalization.

This work provides insight to help future proteomics study designers make more informed decisions on whether physical normalization is necessary for their experiments. This will allow important time and cost saving optimizations to be made to many proteomics workflows.

## Author Contributions

A.Z., G.E.M., H.-Y.Y., C.C.W., and M.J.M. contributed to conceptualization. A.Z., G.E.M., and M.J.M. contributed to experimental design. J.L.I., S.E.C, J.-H.M., K.H.W. and CW contributed to sample production. A.Z., G.E.M. and B.M., contributed to sample preparation. A.Z., and G.E.M. contributed to data collection. A.Z., G.E.M., A.M. and M.R. contributed to data analysis. A.Z. and M.R. contributed to figure generation. M.R. and A.M. contributed to computational normalization software development. The manuscript was written by A.Z. with contributions from all authors. All authors discussed the results and commented on the manuscript. All authors have given approval to the final version of the manuscript.

## Notes

The authors declare the following competing financial interest(s): The MacCoss Lab at the University of Washington has a sponsored research agreement with Thermo Fisher Scientific, the manufacturer of the instrumentation used in this research. M.J.M. is a paid consultant for Thermo Fisher Scientific. The remaining authors declare no competing interests.

## Acknowledgments

We would like to thank Cerise Bennett, Soo Park, William W. Fisher and Richard Weiszmann for their technical assistance with MatTek sample generation. We would also like to thank our colleagues in UW Radiation Oncology and Spectragen Informatics who were part of our UW TEI-REX team.The work presented here was supported by the National Institutes of Health, National Institute of General Medical Sciences under Award P41GM103533 (to M.J.M.). This research is based upon work supported in part by the Office of the Director of National Intelligence (ODNI), Intelligence Advanced Research Projects Activity (IARPA) under Targeted Evaluation of Ionizing Radiation EXposure (TEI-REX) program contract 20008-D2021-2107310008 (LBNL) and through the Army Research Office contract W911NF2220059. The views and conclusions contained should not be interpreted as necessarily representing the official policies, either expressed or implied, of ODNI, IARPA, ARO, or the U.S. Government. The U.S. Government is authorized to reproduce and distribute preprints for governmental purposes notwithstanding any copyright annotation therein.

